# BirdNET can be as good as experts for acoustic bird monitoring in a European city

**DOI:** 10.1101/2024.09.17.613451

**Authors:** Andrew J. Fairbairn, Josija-Simeon Burmeister, Wolfgang W. Weisser, Sebastian T. Meyer

**Author notes:** **Author contributions** AJF and STM developed the research goal and aims. Field data were collected and processed by AJF. AJF analysed the data with input from JSB and STM. The first draft of the manuscript was written by AJF with input from STM. All authors edited and reviewed the final manuscript.

## Abstract

BirdNET has become a leading tool for recognising bird species in audio recordings. However, its applicability in ecological research has been questioned over the sometimes large number of species falsely identified. Using species-specific confidence thresholds has been identified as a powerful approach to solving this issue. However, determining these thresholds is time and resource-consuming. While optimising the parameter setting of the algorithm could be an alternative strategy, the effect of parameter settings on the algorithm’s performance is not well understood. Here, we compared the species identification of BirdNET against expert identification using an acoustic dataset comprising 930 minutes of recordings collected in Munich, Germany. The performance of BirdNET was evaluated using three performance metrics: precision, recall, and F1-score. The metrics were calculated using 24 combinations of the parameters: week, sensitivity, and overlap at four temporal aggregations (pooling of data across time intervals) to also test the effects of recording length. We found that BirdNET closely matched expert identification, particularly when given more data (higher temporal aggregation, F1 score = 0.84) and when including the parameters week of the year, a suitable sensitivity, and an overlap of one to two seconds. Thus, while there are still limitations, using appropriate parameter settings and recording durations, BirdNET yields results comparable to experts without the need for time-consuming estimation of species-specific thresholds. This approach offers reliable presence-absence data in a fast and efficient way while species-specific thresholds are not readily available.

## Introduction

Monitoring bird communities traditionally relies on field observations, requiring experts to manually identify species through visual or auditory cues. While effective, this process is time-intensive, subject to observer bias [1,2], and limited in spatial and temporal coverage [3]. Passive acoustic monitoring (PAM) has emerged as an alternative, allowing continuous data collection across multiple sites simultaneously. However, the bottleneck in analysing the vast amounts of audio data generated through PAM has limited its practical application in ecological research, as manual species identification in recordings remains equally time-consuming and requires specialised expertise [4].

Recent advancements in machine learning have transformed acoustic data analysis, making automated species identification increasingly accessible to researchers without computer science expertise. Among these tools, BirdNET [5] has emerged as a leader tool. The current version, 2.4, has global coverage of over 6,500 avian and non-avian classes [6]. In Germany, BirdNET covers 407 of the 527 bird species tracked by the German Ornithologists Society, including rare and vagrant species [7]. This extensive coverage, coupled with its open-source nature and user-friendly interface, has driven BirdNET’s rapid adoption in both industry applications and scientific research [e.g. 5–7].

Despite its growing popularity, integrating BirdNET in ecological research has been questioned over the sometimes large number of species falsely identified. Using species-specific confidence thresholds has been identified as a powerful approach to solving this issue [6]. As such, many BirdNET studies have focused on optimising confidence thresholds [e.g. 10,11]. However, determining these thresholds is time and resource-consuming. Optimising the parameter setting of the algorithm could be an alternative strategy to improve suboptimal classifications and increase the reliability of ecological metrics derived from BirdNET analyses. Yet the effect of parameter settings—such as overlap, sensitivity, and week of the year—on BirdNET’s performance is not well understood [9], especially in an urban environment.

Here, we used expertly identified acoustic recordings collected in Munich, Germany, to test BirdNET for bird species classification. We aim to determine whether BirdNET can provide species lists comparable to an expert ornithologist in an urban environment. More specifically, we a) assess the impact of varying BirdNET parameters on classification performance, b) examine how different recording length influence output, and c) compare BirdNET’s performance to expert annotations in terms of species richness and identification accuracy. Based on these findings, we provide practical recommendations for parameter settings and validation approaches that optimize BirdNET performance in urban acoustic surveys.

## Methods

### Acoustic recording

We placed a single Frontier Labs BAR on the roof of a housing complex in Munich, Laim, between May and October 2021. We recorded in week-long blocks, with a minimum of one week between recording periods. We recorded one minute every 10 minutes from two hours before sunrise to three hours after sunrise [13] to keep the amount of manual identification required to a manageable level resulting in a total of 15.5 hours of recordings over 60 days. Recordings were taken at a sample rate of 48kHz, a bit depth of 16 and a gain of 40dB.

### Species Identification

Two experts (A. Fairbairn, J.S. Burmeister) identified all bird vocalisations in each recording visually and aurally using Kaleidoscope Pro version 5.6.8 to view the spectrograms and listen to the recordings [14] resulting in a list of species per each one-minute recording. Next, we ran BirdNET analyser v2.3 on the same recordings, producing a list of BirdNET species detections for each one-minute recording.

### Analysis

#### Parameter effects

We tested four parameter settings. First, BirdNET can use the *week* of the year of recording in conjunction with the location to filter what species are likely to occur at a location at that time of the year using eBird [15] species lists. We ran all analyses with and without week included. Second, the detection *sensitivity* (range 0.5 to 1.5, default 1.0) affects how sensitive BirdNET is to faint or background vocalisations. We ran all analyses with three sensitivity levels (0.5, 1.0, 1.5). Third, BirdNET works on three-second audio segments for analysis. The *overlap* determines how many seconds of the previous segment is “overlapped” (default 0.0s). We ran all analyses with four levels of overlap (0, 1, 2, 2.9 seconds). Finally, BirdNET provides a *confidence* level, i.e., the confidence that BirdNET has in its own predictions [6], for each detection. Setting a minimum confidence level in BirdNET causes all detections with a lower confidence to be removed from the results. Thus, to identify the best settings, including the best minimum confidence level, we ran all analyses with the default minimum confidence level (0.1) to get a full list of detections that we could filter afterwards for the analyses.

#### Temporal aggregation

To test how different temporal resolutions (i.e., short versus long recording periods) affect BirdNET’s performance, we aggregated both our reference data and BirdNET results to four different temporal scales: minute (no aggregation), day, week, and the entire dataset (Table 1). For BirdNET results, we first filtered the raw detections based on confidence thresholds and other parameters before aggregation. For each temporal scale, we recorded only species presence data, ignoring repeated detections of the same species. At the minute resolution, if a species was detected multiple times within the same minute, it was recorded as a single presence. At the day resolution, if a species was detected in any minute during that day, it was recorded as a single presence for the entire day. At the week resolution, any species detected at any point during the week was recorded as a single presence for that week.

**Table 1.**
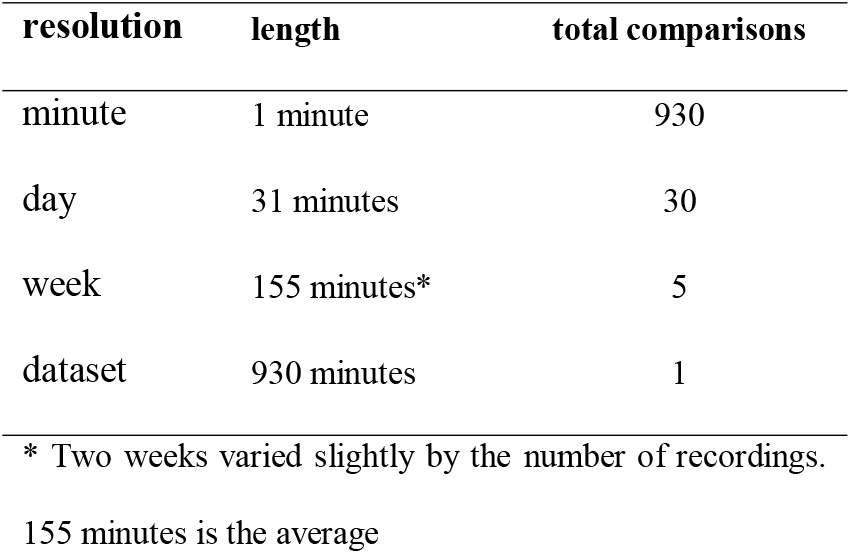
Breakdown of how the data was aggregated to get the different temporal resolutions.

For the entire dataset, each species was recorded as either present or absent overall. This presence-based aggregation was applied to both the expert identifications and filtered BirdNET outputs before making any comparisons. Using the settings that produced the best overall F1 scores, we then calculated separate F1 scores for each individual minute, day, and week to assess how performance variability changes across different temporal resolutions.

#### BirdNET vs expert

To compare BirdNET’s output to expert identifications, we calculated three metrics for each temporal resolution: true positives, defined as species correctly detected as present by BirdNET; false positives, defined as species incorrectly reported as present by BirdNET but not confirmed by experts; and false negatives, defined as species confirmed present by experts but not detected by BirdNET (Fig. 1). We did not calculate true negatives, as there is no meaningful “absence” class in this context—only species that may have been missed by either the expert or BirdNET. Based on these values, we calculated three commonly used machine learning evaluation metrics: precision, recall, and F1 score, to assess BirdNET’s performance. Precision answers the question “How reliable are BirdNET’s identifications?” by measuring the proportion of BirdNET’s species detections that were correct (Eq. 1). A high precision means that when BirdNET identifies a species, it’s likely to be accurate. Recall answers the question “How comprehensive is BirdNET’s coverage?” by measuring the proportion of actually present species (as identified by experts) that BirdNET successfully detected (Eq. 2). A high recall means BirdNET is capturing most of the species present in recordings. Because there is typically a trade-off between precision and recall—improving one may reduce the other—we use the F1 score (Eq. 3) as a balanced performance metric. The F1 score represents the harmonic mean of precision and recall, providing a single value that is high only when both precision and recall are high. This offers an integrated measure of BirdNET’s overall effectiveness at correctly identifying bird species while minimising both false identifications and missed species. Importantly, these metrics were calculated at each temporal resolution, meaning we compared the complete species list for each time unit (minute, day, week, or entire dataset) generated by BirdNET to that identified by the expert rather than evaluating individual detections.

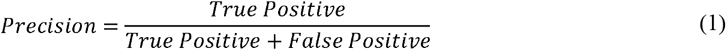

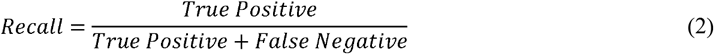

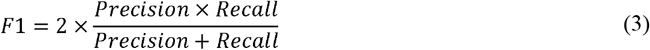

**Figure 1.**
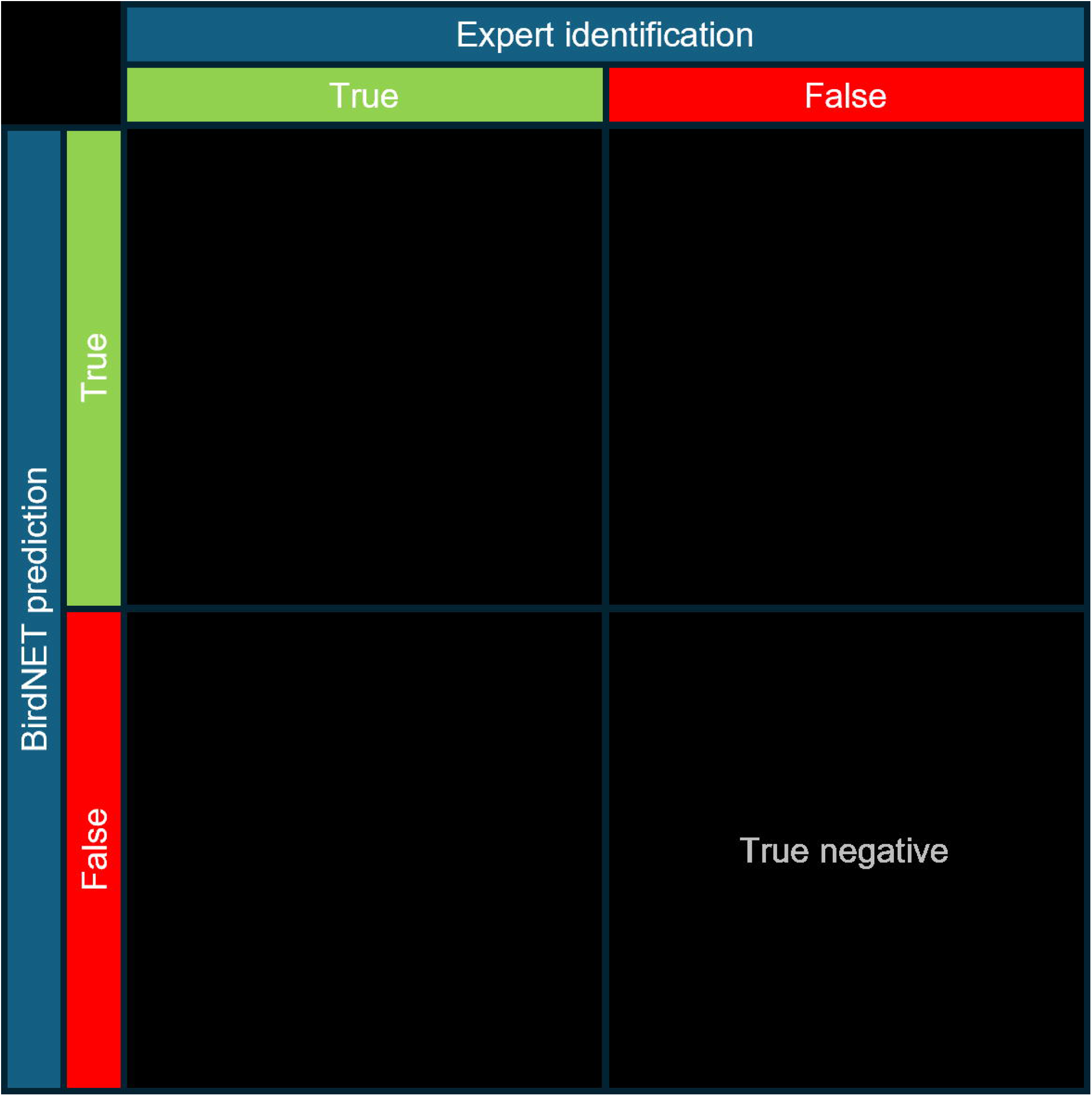
Multiclass confusion matrix showing the definitions of true positive, false negative, and false positive. True negative is light grey because we did not calculate it.

#### Test of parameter settings on BirdNet performance

To understand the effects of each parameter, we fitted a linear mixed-effects model with F1 score as the response variable. Fixed effects included the main effects of aggregation level, sensitivity, overlap, and minimum confidence (minconf^2^), as well as all pairwise interactions among them. A random intercept was included for each run to account for repeated runs of BirdNET. Models were fitted using the *lme* function from the *nlme* package [16] in R [17] (Eq 4; Supplementary Table S1).

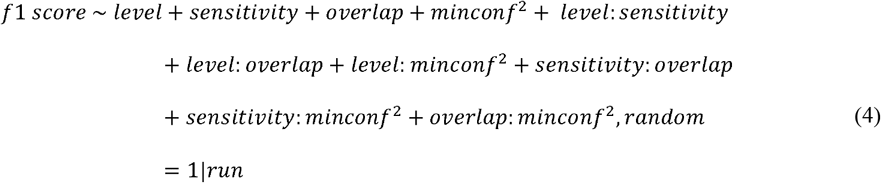

#### Confirmation test

While in the previous steps, we assume that the expert identification is perfect, incorrect identification or missed species may also occur in the species lists based on expert identification, despite our best efforts. Therefore, we conducted a confirmation test assuming errors can occur within expert and BirdNET identification. Using the parameter values that produced the best result for the dataset resolution (highest F1 score), we manually checked a portion of the BirdNET detections. Following Sethi et. al. 2021 [18], we sorted the BirdNET results by species and randomly selected up to 50 results for each species. For species with fewer than 50 detections, we reviewed all available detections. Each selected detection was re-examined by listening to the audio and confirming whether it was correctly identified. We additionally examined the confidence ranges of any false negative species (species identified by the experts but missed by BirdNET when using the best parameter settings) to determine what the impact of lowering the confidence threshold would be.

## Results

We recorded 930 minutes over five weeks and expertly identified 9466 vocalisations of a total of 23 species in the recordings. The most commonly identified species were Short-toed Treecreeper *Certhia brachydactyla* (n=2894), Blackbird *Turdus merula* (n=1078), Common Swift *Apus apus* (n=817) and Great Tit *Parus major* (n=798; Supplementary Table S2). With default settings, and including week of the year, BirdNET identified 93 species with the most frequently identified being the European Robin *Erithacus rubecula* (n=2890), the Blackbird (n=1820), the Great Tit (n=1264), and the Short-toed Treecreeper (n=1037; Supplementary Table S3).

### BirdNET vs expert

When comparing BirdNET with expert identification, temporal aggregation and all tested parameters (minimum confidence, week, sensitivity, and overlap) significantly affected BirdNET performance (F1 scores; Fig. 2, Supplementary Table S4). Including the *week* of the year consistently provided better results (Supplement Fig. S1). Minimum confidence had the strongest effect on F1 score (p < 0.001) and significantly interacted with temporal aggregation, sensitivity, and overlap (Fig. 2, Supplement Table S1). As aggregation increased from minute resolution to the whole dataset, maximum F1 scores improved from 0.62 to 0.84, respectively, when using optimal, i.e. those providing the best F1 scores, combinations of minimum confidence, sensitivity, and overlap (Fig. 2, Table 2). With the exception of high confidence levels for the minute and day aggregations, decreasing overlap provided higher maximum F1 scores (Fig. 2, Supplement Table S4). Default BirdNET parameter settings performed poorly when considering F1 scores (Table 2). However, default settings provided higher recall, maximising the number of true positives while inflating false positives.

**Table 2.**
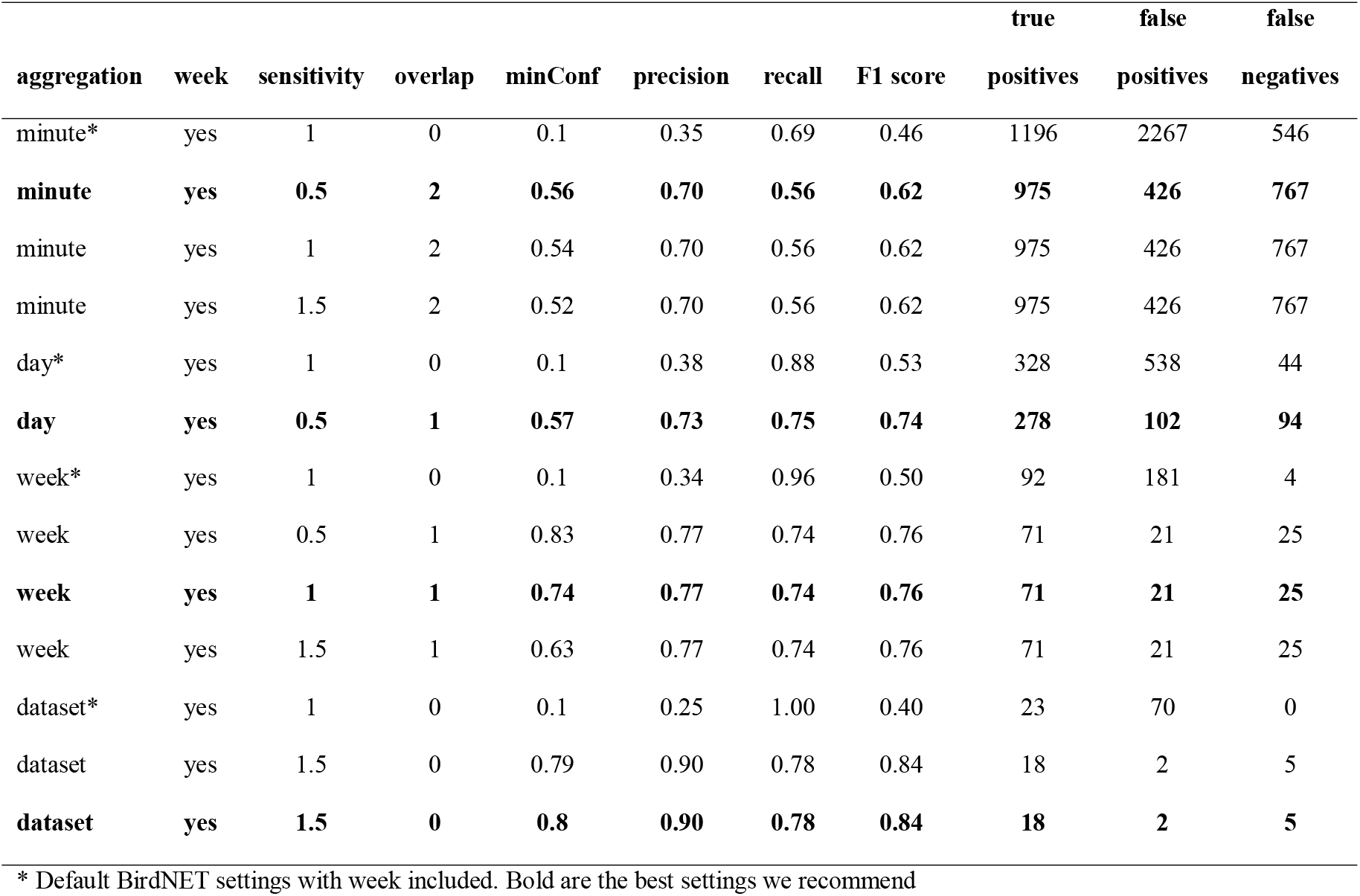
Best settings resulted in the highest F1 scores for each temporal aggregation level from 1,944 BirdNET and expert.

**Figure 2.**
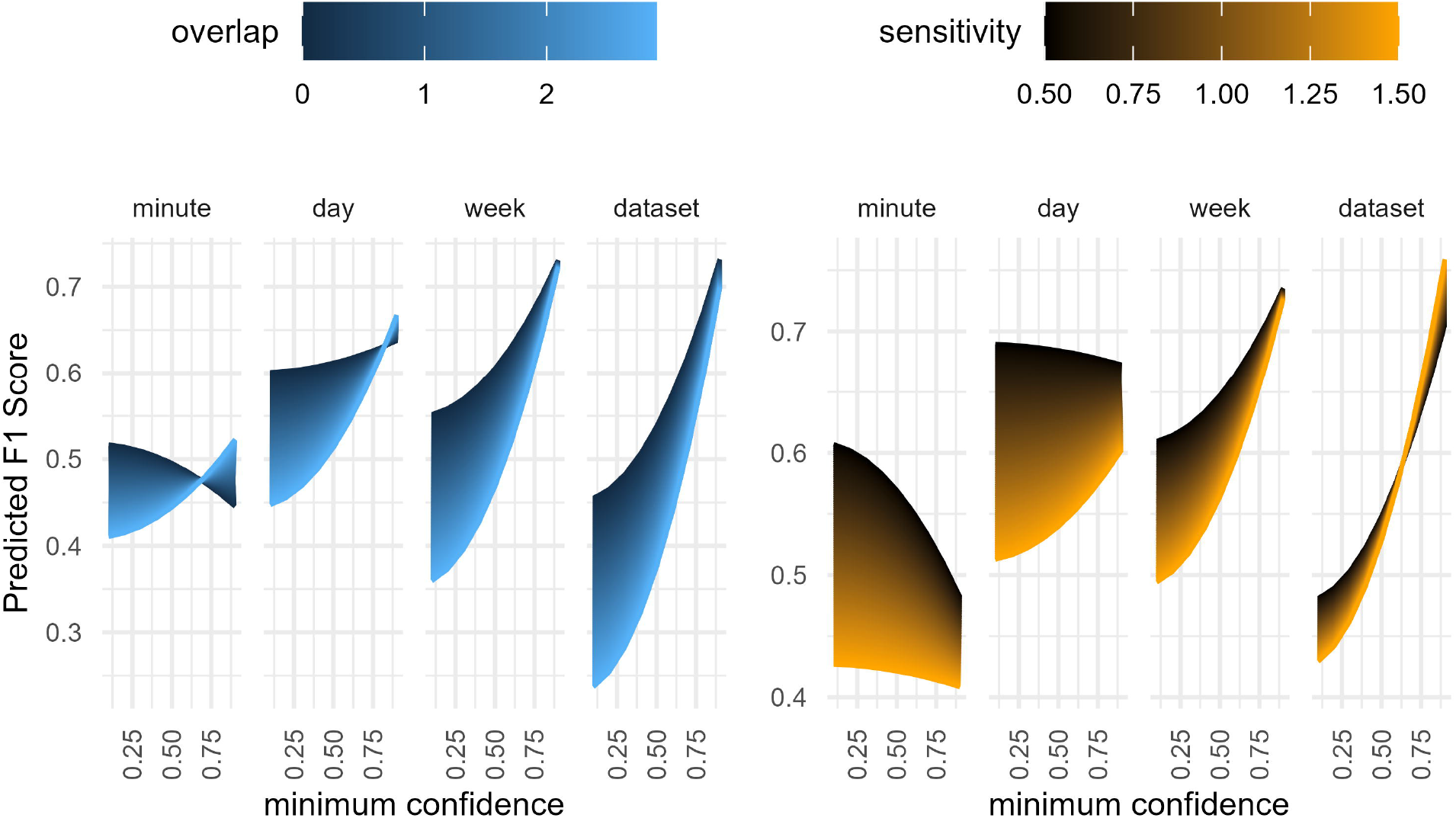
Predicted F1 score as a function of data aggregation level, overlap, sensitivity and minimum confidence from a liner mixed-effects model based on 1,944 comparison tests between BirdNET and an expert ornithologist. Each panel shows the variation in predicted F1 scores across each aggregation level, minimum confidence and overlap (left) or sensitivity (right).

### Confirmation test

In our confirmation test, conducted using the best parameter settings identified for data aggregated across the entire dataset, BirdNET detected 20 species. Among these were two species—*Delichon urbicum* (Western house martin) and *Turdus philomelos* (Song thrush)—that were not present in the expert identifications and were flagged as false positives. However, after manually reviewing the audio clips associated with these detections, we confirmed that all 14 detections of Song thrush were, in fact, correct. Only the single detection of Western house martin remained a true false positive (Fig. 3). With these adjustments, the F1 score for the full-dataset resolution was revised to 0.86 (Precision = 0.95, Recall = 0.792). The expert identification included an additional five species missed by BirdNET. It is important to note that these false negatives were not addressed in the confirmation test, as they represent species that were not detected by the model. In our check of the false negative species, we found that their confidence scores ranged from a max of 0.37 to a max of 0.68 (Supplementary Table S5). As such, a lower minimum confidence threshold would have included some of these species, but at the cost of additional false positives. Lowering the minimum confidence to 0.54, for example, reduced the false negatives to 2, but at the cost of an additional 9 false positive species, which would result in an overall lower F1 score.

**Figure 3.**
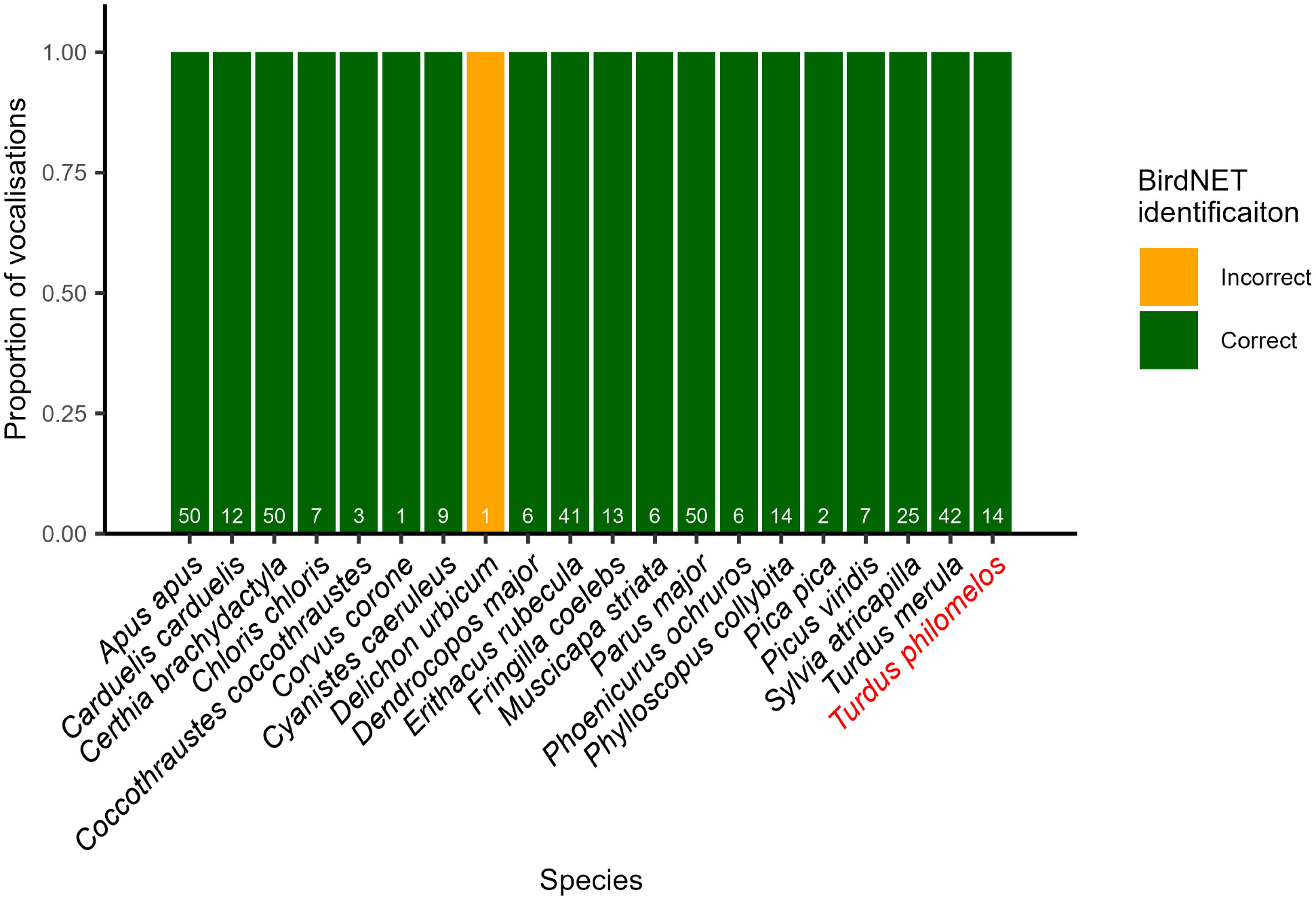
The proportion of BirdNET detections from the dataset resolution that were manually checked by an expert and determined to be correct or incorrect. A maximum of 50 random detections per species were checked. If a species was detected less than 50 times, all detections were checked. White numbers are the number of detections checked (if less than 50, only that number were available). The red label denotes the species missed in our expert identification that BirdNET identified.

## Discussion

Our study adds to the growing body of work evaluating the performance of BirdNET and is to our knowledge one of the first to systematically investigate the effect of varying BirdNET parameters. While current acoustic monitoring practices recommend short recordings during peak activity periods [13,19], BirdNET eliminates the need to listen to entire recordings, allowing for longer and more frequent data collection. As highlighted by our temporal resolution results and as others have found [20], BirdNET performs better with longer recordings, which also enhances monitoring by capturing species with varying activity patterns [21]. Adjusting the parameters, more specifically, by including the week of the year, increasing the overlap to one or two seconds, and using a higher than default confidence threshold produced better results, especially when using short recordings as we did here. When provided with the correct settings and sufficient data (e.g., aggregated over longer periods or longer recordings), BirdNET performed nearly as well as an expert and, in our case, even detected a species missed by the expert. However, it did miss five species identified by the experts. Nevertheless, we show that BirdNET can be used to monitor birds in acoustically complex environments such as cities.

While the best settings for BirdNET parameters varied depending on the temporal resolution level, we can deduce some generalisations for running BirdNET. Since our best-performing parameter settings always included the week of the year, we recommend general monitoring to include the week of the year and location. Overlap had a more significant impact on performance than sensitivity. We recommend using an overlap between one and two seconds for short one to five-minute recording schemes. With longer or continuous recordings, an overlap may not be necessary as vocalisations that may be missed or unidentifiable from being cut are likely to occur again. Default sensitivity (1.0) generally produced the same results as higher or lower sensitivities while maintaining higher minimum confidence levels. We, therefore, recommend using the default sensitivity. For short recordings of 30 minutes or less, we suggest filtering by a minimum confidence of 0.54 or higher. For longer recordings, a higher minimum confidence yields the most reliable results, although such a high threshold may exclude some low-confidence but valid detections.

Despite its performance, we recommend that the results of BirdNET be validated, especially when using very short recordings. While research goals will dictate the amount of validation necessary, we recommend a few quick methods to ensure the best results. If the researcher is familiar with what is likely to occur on their study site, only manually checking unlikely or uncommon species is likely to suffice. If confirming the presence of the species at a location (i.e. producing a species list) is the research goal, validation can be done easily by manually checking the top results for each species/site, as only one valid detection is needed to confirm occurrence [9]. Additionally, removing singletons or doubletons and checking only infrequently detected species is likely to provide more accurate species lists than just the raw output of BirdNET. As our confirmation test showed, had we lowered the confidence level to 0.54, removed the singletons and checked the species that were detected 10 or fewer times, we would have had a species list closer to that of the expert, missing only two species. This highlights how simple validation and filtering steps can significantly improve agreement between expert and automated methods.

We recognise that current practice recommends using individual species thresholds as model performance can vary greatly between species [6,12]. We think it is important to let the research question dictate which method should be used. Creating individual species scores requires a significant upfront investment in time and requires expert knowledge of the different vocalisations a species can make. Further, the creation of species-specific thresholds assumes that a dataset contains enough detections across the confidence range, although a universal number of detections has yet to be determined. Research questions for which the number of valid vocalisations is important, e.g. studies investigating activity patterns [10], could benefit from species-specific thresholds as they maintain a greater number of detections. For example, Tseng et al. (2025) [12] found that using individual species scores retained a much larger number of detections (70 ± 37%) than a universal threshold (17 ± 14%) as we use here. Still, it has yet to be determined if these thresholds are transferrable across time (e.g., season, time of day, years) and space (e.g., regions and habitats). Until standardised species-specific thresholds become available, for short-term studies or rapid biodiversity assessments, universal thresholds with simple validation procedures may be sufficient and more resource-efficient. Therefore, it is important to consider the aims of a project when deciding if a universal threshold is adequate or if individual thresholds should be calculated.

Our study provides additional support for BirdNET as a practical tool for species identification, particularly in urban environments, but also highlights some limitations that require careful consideration. We show that appropriate recording strategies and utilising or adjusting key parameters—such as including week of the year, increasing overlap for short recordings, and using a higher minimum confidence threshold—can substantially improve detection performance. It should be noted that our results represent a single site in a southern German city and results from different regions or environments may vary. While we acknowledge that the universal confidence thresholds we propose may not suit all research contexts, with basic validation or filtering (e.g., checking uncommon or infrequent species), they can still yield ecologically useful results. The universal threshold approach offers reliable presence-absence data in a fast and efficient way as long as species-specific thresholds are not readily available. BirdNET effectively overcomes some of the limitations of conventional ornithological sampling methods, thus positioning it as a valuable asset in the ongoing quest for comprehensive and efficient biodiversity monitoring practices, offering new research opportunities in ecology and ornithology.

## Acknowledgements

We thank the German Research Foundation (DFG Research Training Group 2679 - Urban Green Infrastructure) for funding this research. We would also like to thank Julia Windl and Lisa Maier for assisting with identifying the bird recordings. Additionally, we would like to thank Münchener Wohnen for permitting our recording.

## Notes

### Competing Interest Statement

The authors have declared no competing interest.

### Summary of Updates

Expanded and updated introduction. Expanded and updated discussion. Clarified methods.

## References

1. Scher CL, Clark JS. Species traits and observer behaviors that bias data assimilation and how to accommodate them. Ecological Applications. 2023;33: e2815. doi:10.1002/eap.2815

2. Harris JBC, Haskell DG. Simulated Birdwatchers’ Playback Affects the Behavior of Two Tropical Birds. PLOS ONE. 2013;8: e77902. doi:10.1371/journal.pone.0077902

3. Kułaga K, Budka M. Bird species detection by an observer and an autonomous sound recorder in two different environments: Forest and farmland. Pérez-García JM, editor. PLoS ONE. 2019;14: e0211970. doi:10.1371/journal.pone.0211970

4. Hoefer S, McKnight, Donald T., Allen-Ankins, Slade, Nordberg, Eric J., and Schwarzkopf L. Passive acoustic monitoring in terrestrial vertebrates: a review. Bioacoustics. 2023;32: 506–531. doi:10.1080/09524622.2023.2209052

5. Kahl S, Wood CM, Eibl M, Klinck H. BirdNET: A deep learning solution for avian diversity monitoring. Ecol Inform. 2021;61. doi:10.1016/j.ecoinf.2021.101236

6. Wood CM, Kahl S. Guidelines for appropriate use of BirdNET scores and other detector outputs. J Ornithol. 2024;165: 777–782. doi:10.1007/s10336-024-02144-5

7. Barthel PH, Krüger T. Liste der Vögel Deutschlands: Version 3.2. Deutsche Ornithologen-Gesellschaft e.V.; 2019. Available: http://www.do-g.de/fileadmin/BarthelKrueger_2019_Liste_der_Voegel_Deutschlands_3.2_DO-G.pdf

8. Sethi SS, Fossøy F, Cretois B, Rosten CM. Management relevant applications of acoustic monitoring for Norwegian nature – The Sound of Norway. Norsk institutt for naturforskning (NINA); 2021. Available: https://hdl.handle.net/11250/2832294

9. PérezLGranados C. BirdNET: applications, performance, pitfalls and future opportunities. Ibis. 2023;165: 1068–1075. doi:10.1111/ibi.13193

10. Amorós-Ausina D, Schuchmann K-L, Marques MI, Pérez-Granados C. Living Together, Singing Together: Revealing Similar Patterns of Vocal Activity in Two Tropical Songbirds Applying BirdNET. Sensors. 2024;24: 5780. doi:10.3390/s24175780

11. David Funosas, Luc Barbaro, Laura Schillé, Arnaud Elger, Bastien Castagneyrol, Maxime Cauchoix. Assessing the potential of BirdNET to infer European bird communities from large-scale ecoacoustic data. bioRxiv. 2023; 2023.12.06.570351. doi:10.1101/2023.12.06.570351

12. Tseng S, Hodder DP, Otter KA. Setting BirdNET confidence thresholds: species-specific vs. universal approaches. J Ornithol. 2025 [cited 7 Apr 2025]. doi:10.1007/s10336-025-02260-w

13. Abrahams C. Bird bioacoustic surveys – Developing a standard protocol. In Practice. 2018: 20–23.

14. Wildlife Acoustics. Kaleidoscope Pro 5. Wildlife Acoustics; 2021. Available: https://www.wildlifeacoustics.com/products/kaleidoscope-pro

15. Sullivan BL, Wood CL, Iliff MJ, Bonney RE, Fink D, Kelling S. eBird: A citizen-based bird observation network in the biological sciences. Biological Conservation. 2009;142: 2282–2292. doi:10.1016/j.biocon.2009.05.006

16. Pinheiro J, Bates D, DebRoy S, Sarkar D, R Core Team. nlme: Linear and nonlinear mixed effects models. 2021. Available: https://CRAN.R-project.org/package=nlme

17. R Core Team. R: A Language and Environment for Statistical Computing. Vienna, Austria: R Foundation for Statistical Computing; 2023. Available: https://www.R-project.org/

18. Sethi SS, Fossøy, F, Cretois, B, Rosten, C. M. Management relevant applications of acoustic monitoring for Norwegian nature – The Sound of Norway. Norwegian Institute for Nature Research; 2021 p. 37. Report No.: NINA Report 2064.

19. Metcalf O, Abrahams C, Ashington B, Baker E, Bradfer-Lawrence T, Browning E, et al. Good practice guidelines for long-term ecoacoustic monitoring in the UK. The UK Acoustics Network; 2023 Feb pp. 1–82. Available: https://acoustics.ac.uk/

20. Cole JS, Michel NL, Emerson SA, Siegel RB. Automated bird sound classifications of long-duration recordings produce occupancy model outputs similar to manually annotated data. Ornithol Appl. 2022;124: 1–15. doi:10.1093/ornithapp/duac003

21. Robbins CS. Effect of time of day on bird activity. C. John Ralph, J. Michael Scott, editors. Studies in Avian Biology. 1981;6: 275–286.

