## Supplementary information for "BirdNET can be as good as experts for acoustic bird monitoring in a European city"

| Supplementary Table S1. ANOVA results from a linear mixed effects model for F1 score and the BirdNET parameters sensitivity, overlap, and minimum confidence plus four data aggregation levels. | |
| --- | --- |
| **variable** | **F and p value** |
| level | F_3,63_=147.21; p<0.001 |
| sensitivity | F_1,20_=5.11; p=0.035 |
| overlap | F_1,20_=9.83; p=0.005 |
| minconf^2 | F_1,7674_=5023.03; p<0.001 |
| level:sensitivity | F_3,63_=35.17; p=0.001 |
| level:overlap | F_3,63_=19.56; p=0.001 |
| level:minconf^2 | F_3,7674_=750.56; p<0.001 |
| sensitivity:overlap | F_1,20_=0.43; p=0.521 |
| sensitivity:minconf^2 | F_1,7674_=239.59; p<0.001 |
| overlap:minconf^2 | F_1,7674_=603.84; p<0.001 |

| Supplementary Table S2. Expertly identified species with the total number of times each species was identified in the data. | |
| --- | --- |
| **Species** | **n identifications** |
| *Certhia brachydactyla* | 2894 |
| *Turdus merula* | 1078 |
| *Apus apus* | 817 |
| *Parus major* | 798 |
| *Sylvia atricapilla* | 555 |
| *Erithacus rubecula* | 527 |
| *Corvus corone* | 458 |
| *Phylloscopus collybita* | 401 |
| *Cyanistes caeruleus* | 380 |
| *Fringilla coelebs* | 323 |
| *Chloris chloris* | 321 |
| *Muscicapa striata* | 263 |
| *Phoenicurus ochruros* | 179 |
| *Passer montanus* | 136 |
| *Carduelis carduelis* | 118 |
| *Dendrocopos major* | 82 |
| *Coccothraustes coccothraustes* | 40 |
| *Picus viridis* | 28 |
| *Sitta europaea* | 26 |
| *Columba palumbus* | 15 |
| *Regulus regulus* | 12 |
| *Pica pica* | 8 |
| *Streptopelia decaocto* | 7 |
| **Total** | 9466 |

| Supplementary Table S3. Species identified by BirdNET (default settings, minimum confidence 0.1) with the total number of times each species was identified. | |
| --- | --- |
| **Species** | **n identifications** |
| *Erithacus rubecula* | 2890 |
| *Turdus merula* | 1820 |
| *Parus major* | 1264 |
| *Certhia brachydactyla* | 1037 |
| *Apus apus* | 893 |
| *Turdus philomelos* | 868 |
| *Corvus corone* | 622 |
| *Sylvia atricapilla* | 532 |
| *Cyanistes caeruleus* | 462 |
| *Phylloscopus collybita* | 326 |
| *Carduelis carduelis* | 292 |
| *Chloris chloris* | 267 |
| *Coccothraustes coccothraustes* | 245 |
| *Fringilla coelebs* | 242 |
| *Passer domesticus* | 242 |
| *Anthus trivialis* | 207 |
| *Muscicapa striata* | 196 |
| *Phoenicurus ochruros* | 163 |
| *Corvus cornix* | 152 |
| *Dendrocopos major* | 122 |
| *Delichon urbicum* | 104 |
| *Motacilla cinerea* | 77 |
| *Motacilla alba* | 52 |
| *Actitis hypoleucos* | 48 |
| *Fulica atra* | 48 |
| *Regulus regulus* | 44 |
| *Turdus viscivorus* | 36 |
| *Picus viridis* | 29 |
| *Columba palumbus* | 28 |
| *Gallinula chloropus* | 24 |
| *Chroicocephalus ridibundus* | 23 |
| *Emberiza schoeniclus* | 23 |
| *Prunella modularis* | 23 |
| *Periparus ater* | 21 |
| *Aegithalos caudatus* | 19 |
| *Regulus ignicapilla* | 19 |
| *Charadrius dubius* | 18 |
| *Corvus frugilegus* | 18 |
| *Alauda arvensis* | 17 |
| *Pica pica* | 17 |
| *Emberiza citrinella* | 16 |
| *Troglodytes troglodytes* | 16 |
| *Ficedula hypoleuca* | 15 |
| *Motacilla flava* | 15 |
| *Poecile palustris* | 15 |
| *Anas platyrhynchos* | 14 |
| *Anas crecca* | 12 |
| *Turdus pilaris* | 11 |
| *Ardea cinerea* | 10 |
| *Linaria cannabina* | 10 |
| *Buteo buteo* | 8 |
| *Alcedo atthis* | 6 |
| *Anthus pratensis* | 5 |
| *Corvus corax* | 5 |
| *Dryocopus martius* | 5 |
| *Emberiza calandra* | 5 |
| *Passer montanus* | 5 |
| *Sitta europaea* | 5 |
| *Falco tinnunculus* | 4 |
| *Grus grus* | 4 |
| *Phoenicurus phoenicurus* | 4 |
| *Pyrrhula pyrrhula* | 4 |
| *Rallus aquaticus* | 4 |
| *Serinus serinus* | 4 |
| *Spinus spinus* | 4 |
| *Columba livia* | 3 |
| *Mareca strepera* | 3 |
| *Oriolus oriolus* | 3 |
| *Saxicola rubetra* | 3 |
| *Saxicola rubicola* | 3 |
| *Streptopelia decaocto* | 3 |
| *Alopochen aegyptiaca* | 2 |
| *Corvus monedula* | 2 |
| *Fringilla montifringilla* | 2 |
| *Hirundo rustica* | 2 |
| *Lophophanes cristatus* | 2 |
| *Loxia curvirostra* | 2 |
| *Milvus milvus* | 2 |
| *Poecile montanus* | 2 |
| *Sturnus vulgaris* | 2 |
| *Sylvia borin* | 2 |
| *Tringa nebularia* | 2 |
| *Anser anser* | 1 |
| *Bucephala clangula* | 1 |
| *Certhia familiaris* | 1 |
| *Circus aeruginosus* | 1 |
| *Curruca communis* | 1 |
| *Curruca curruca* | 1 |
| *Luscinia megarhynchos* | 1 |
| *Sterna hirundo* | 1 |
| *Tachybaptus ruficollis* | 1 |
| *Tringa ochropus* | 1 |
| *Vanellus vanellus* | 1 |

| Supplementary Table S4. Summary statistics of F1 score as it varies with minimum confidence level. | | | | | | | |
| --- | --- | --- | --- | --- | --- | --- | --- |
| **resolution** | **overlap** | **sensitivity** | **min F1** | **max F1** | **mean F1** | **median F1** | **sd F1** |
| minute | 0 | 0.5 | 0.42 | 0.58 | 0.54 | 0.56 | 0.04 |
| minute | 0 | 1 | 0.33 | 0.58 | 0.51 | 0.54 | 0.07 |
| minute | 0 | 1.5 | 0.18 | 0.58 | 0.42 | 0.45 | 0.13 |
| minute | 1 | 0.5 | 0.45 | 0.60 | 0.56 | 0.58 | 0.04 |
| minute | 1 | 1 | 0.36 | 0.60 | 0.53 | 0.55 | 0.06 |
| minute | 1 | 1.5 | 0.16 | 0.60 | 0.44 | 0.46 | 0.13 |
| minute | 2 | 0.5 | 0.48 | 0.62 | 0.58 | 0.58 | 0.04 |
| minute | 2 | 1 | 0.39 | 0.62 | 0.54 | 0.56 | 0.06 |
| minute | 2 | 1.5 | 0.14 | 0.62 | 0.44 | 0.46 | 0.14 |
| minute | 2.9 | 0.5 | 0.35 | 0.59 | 0.51 | 0.53 | 0.07 |
| minute | 2.9 | 1 | 0.26 | 0.59 | 0.49 | 0.52 | 0.10 |
| minute | 2.9 | 1.5 | 0.09 | 0.59 | 0.41 | 0.44 | 0.15 |
| day | 0 | 0.5 | 0.61 | 0.71 | 0.68 | 0.68 | 0.03 |
| day | 0 | 1 | 0.50 | 0.71 | 0.65 | 0.67 | 0.05 |
| day | 0 | 1.5 | 0.28 | 0.70 | 0.55 | 0.59 | 0.13 |
| day | 1 | 0.5 | 0.60 | 0.74 | 0.69 | 0.70 | 0.03 |
| day | 1 | 1 | 0.51 | 0.74 | 0.66 | 0.68 | 0.06 |
| day | 1 | 1.5 | 0.29 | 0.74 | 0.56 | 0.58 | 0.14 |
| day | 2 | 0.5 | 0.54 | 0.74 | 0.68 | 0.70 | 0.05 |
| day | 2 | 1 | 0.45 | 0.74 | 0.66 | 0.68 | 0.07 |
| day | 2 | 1.5 | 0.27 | 0.74 | 0.56 | 0.58 | 0.14 |
| day | 2.9 | 0.5 | 0.41 | 0.72 | 0.58 | 0.59 | 0.08 |
| day | 2.9 | 1 | 0.35 | 0.72 | 0.57 | 0.59 | 0.11 |
| day | 2.9 | 1.5 | 0.24 | 0.72 | 0.51 | 0.53 | 0.15 |
| week | 0 | 0.5 | 0.57 | 0.74 | 0.68 | 0.69 | 0.05 |
| week | 0 | 1 | 0.50 | 0.74 | 0.66 | 0.68 | 0.07 |
| week | 0 | 1.5 | 0.36 | 0.73 | 0.60 | 0.62 | 0.10 |
| week | 1 | 0.5 | 0.55 | 0.76 | 0.68 | 0.69 | 0.05 |
| week | 1 | 1 | 0.48 | 0.76 | 0.66 | 0.68 | 0.07 |
| week | 1 | 1.5 | 0.35 | 0.76 | 0.60 | 0.63 | 0.11 |
| week | 2 | 0.5 | 0.50 | 0.74 | 0.64 | 0.66 | 0.06 |
| week | 2 | 1 | 0.45 | 0.74 | 0.63 | 0.66 | 0.08 |
| week | 2 | 1.5 | 0.35 | 0.74 | 0.58 | 0.62 | 0.12 |
| week | 2.9 | 0.5 | 0.43 | 0.70 | 0.54 | 0.53 | 0.06 |
| week | 2.9 | 1 | 0.40 | 0.72 | 0.54 | 0.53 | 0.09 |
| week | 2.9 | 1.5 | 0.33 | 0.72 | 0.53 | 0.53 | 0.13 |
| dataset | 0 | 0.5 | 0.45 | 0.75 | 0.60 | 0.60 | 0.09 |
| dataset | 0 | 1 | 0.40 | 0.75 | 0.59 | 0.60 | 0.11 |
| dataset | 0 | 1.5 | 0.32 | 0.84 | 0.59 | 0.60 | 0.17 |
| dataset | 1 | 0.5 | 0.44 | 0.74 | 0.60 | 0.59 | 0.08 |
| dataset | 1 | 1 | 0.39 | 0.72 | 0.59 | 0.59 | 0.10 |
| dataset | 1 | 1.5 | 0.32 | 0.82 | 0.58 | 0.59 | 0.16 |
| dataset | 2 | 0.5 | 0.41 | 0.67 | 0.52 | 0.53 | 0.07 |
| dataset | 2 | 1 | 0.38 | 0.70 | 0.53 | 0.53 | 0.09 |
| dataset | 2 | 1.5 | 0.31 | 0.83 | 0.54 | 0.53 | 0.16 |
| dataset | 2.9 | 0.5 | 0.36 | 0.56 | 0.44 | 0.44 | 0.05 |
| dataset | 2.9 | 1 | 0.33 | 0.59 | 0.44 | 0.44 | 0.07 |
| dataset | 2.9 | 1.5 | 0.30 | 0.82 | 0.47 | 0.44 | 0.14 |

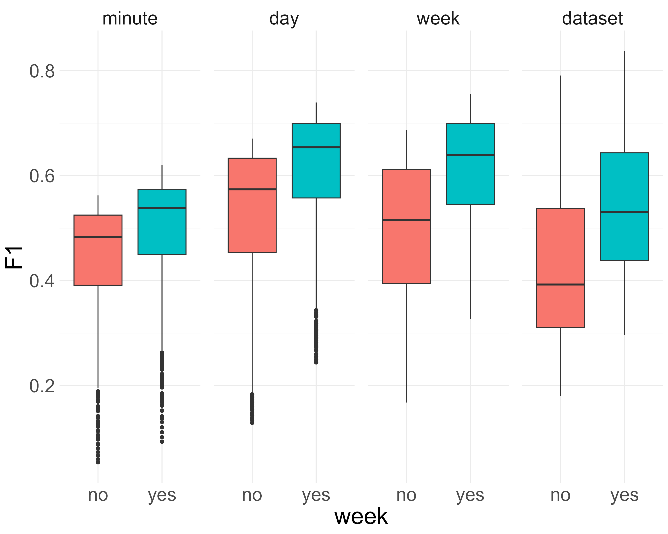

Supplementary Figure S1. Differences in F1 scores for each of the four temporal resolutions when including or not including the week of the year.

| **Supplementary Table S5.** Confidence range of false negative species from the best settings at the dataset resolution. | |
| --- | --- |
| **Species** | **Confidence range** |
| Passer montanus | 0.1-0.42 |
| Sitta europaea | 0.1-0.69 |
| Columba palumbus | 0.1-0.59 |
| Regulus regulus | 0.1-0.55 |
| Streptopelia decaocto | 0.1-0.38 |
